# A unique multi-synaptic mechanism regulates dopamine release in the nucleus accumbens during adolescence

**DOI:** 10.1101/2020.09.18.303974

**Authors:** Elizabeth G. Pitts, Taylor A. Stowe, Mark J. Ferris

## Abstract

Adolescence is characterized by changes in reward-related behaviors, social behaviors, and decision making. These behavioral changes are necessary for the transition into adulthood, but they also increase vulnerability to the development of a range of psychiatric disorders. Major reorganization of the dopamine system during adolescence is thought to underlie, in part, the behavioral changes and increased vulnerability. Here, we utilized fast scan cyclic voltammetry to examine differences in regulation of dopamine release in the nucleus accumbens core of adolescent and adult male rats. We found that differences between adolescent and adult stimulated dopamine release is driven by a unique multisynaptic mechanism in early adolescence involving acetylcholine acting at α6-containing nicotinic acetylcholine receptors to mediate inhibition of dopamine via GABA release. These changes in dopamine regulation across adolescence provides a springboard for our understanding of basic brain development and targeted therapy for a range of psychiatric conditions that emerge in adolescence.

## Introduction

Adolescence is the transitional period between childhood and adulthood and is characterized by changes in behaviors, such as increases in risk-taking, sensation seeking, and peer-focused sociality (Spear, 2000, 2011, 2013; Nelson et al., 2005). These changes in behavior are thought to be evolutionarily advantageous, as they allow for the exploration necessary for a transition to independence (Spear, 2000). However, adolescence is also a time of increased vulnerability to the development of a range of psychiatric disorders, including substance use disorders (SUD), schizophrenia, and affective disorders, that can persist into adulthood (Casey et al., 2008; Fareri et al., 2008; Steinberg, 2008). Adolescent onset is also associated with increased severity of a range of psychiatric disorders (Chambers et al., 2003; Kyriakopoulos and Frangou, 2007; Andersen and Teicher, 2008). This may be in part because the brain is still developing during adolescence and disturbances, such as drug-taking or stress, may disrupt the natural developmental trajectory. Thus, understanding how the system develops in a healthy system may help us understand how and why the system is perturbed in disease.

Reorganization of the dopamine system is hypothesized to underlie many of the changes in behavior, particularly in reward-related learning and decision making, and the increased vulnerability to psychiatric disorders that are common in adolescence (Spear, 2000, 2013; Nelson et al., 2005; Wahlstrom et al., 2010a,b). Preclinical research indicates that the dopamine system is going through major changes in adolescence. For example, the density of D_1_ and D_2_ dopamine receptors in the striatum, firing rate of dopamine neurons, and dopamine synthesis all peak during adolescence (Teicher et al., 1995; Tarazi et al., 1999; Philpot et al., 2009; McCutcheon et al., 2012). While reports examining extracellular levels of dopamine in the striatum show both lower and higher levels of dopamine during adolescence (Badanich et al., 2006; Cao et al., 2007; Philpot et al., 2009), sub-second stimulated dopamine release is consistently found to be decreased in the striatum of adolescent rats (Stamford, 1989; Palm and Nylander, 2014; Pitts et al., 2020). We have previously shown that stimulated dopamine release is decreased in the nucleus accumbens (NAc) core of early adolescent male rats (Pitts et al., 2020), yet, what mechanism is differentially mediating dopamine release in adolescent and adult rats is still unclear.

Since previous research from our lab showed that the NAc core has the largest difference in stimulated dopamine release between adult and adolescent male rats (Pitts et al., 2020), we focused this report on understanding the mechanisms mediating dopamine release in that region. This is of particular importance because the NAc core is important in regulating reward-related learning and goal-directed decision making (Di Chiara, 2002; Saddoris et al., 2013), which are altered in adolescence. Terminal dopamine release in the striatum does not always correspond with action potentials from ventral tegmental and substantia nigra projection neurons (Trulson, 1985; Mohebi et al., 2019). Release is also regulated by local circuitry, by both homosynaptic and heterosynaptic mechanisms (see Nolan et al., 2020). Functionality of dopamine transporters and release probability (homosynaptic regulation) can both mediate stimulated dopamine release (see Ferris et al., 2013). Heterosynaptic mechanisms include neurotransmitter release from local interneurons and projection neurons, such as acetylcholine interneurons, which play a major role in regulating local terminal dopamine release (see Cachope and Cheer, 2014; Collins and Saunders, 2020; Nolan et al., 2020). For example, optogenetic stimulation of acetylcholine release drives terminal dopamine release independent of action potential through activation of nicotinic acetylcholine receptors (nAChRs) on dopamine terminals (Threlfell et al., 2012; Cachope et al., 2012). This local control over terminal dopamine release has important implications for how dopamine encodes salient information and drives behavior, and one or more of the regulatory mechanisms that mediate terminal dopamine release may change between adolescence and adulthood. Here, we use *ex vivo* fast scan cyclic voltammetry (FSCV) to compare stimulated dopamine release and its local circuitry regulation in the NAc core of adult and adolescent male rats. Greater understanding of the developmental changes in dopamine regulation may shed light on mechanisms driving changes in reward-related behaviors, and related increase in vulnerability, that are characteristic of adolescence.

## Results

### Adolescent rats have decreased dopamine release in the NAc core

We first compared stimulated dopamine release in the NAc core of adult (>P70) and early adolescent (P28-35) rats using *ex vivo* FSCV (Figure 1A). Adolescent rats have significantly lower dopamine release to single-pulse stimulations (*t*_15_=2.938, *p*=0.010)(Figure 1B) and across a range of stimulation parameters that model tonic-(5-10 Hz) and phasic-like (20-100 Hz) firing of dopamine neurons (main effect of age: *F*_1, 20_=10.96, *p*=0.004)(Figure 1C). Differences in dynamics at the dopamine terminal can modulate stimulated dopamine release (see Ferris et al., 2013; Nolan et al., 2020). For example, activity of dopamine transporters (DAT) have been shown to impact stimulated dopamine levels (Condon et al., 2019). To determine whether differences in DAT activity may mediate the age-related differences in stimulated dopamine release, we examined *Vmax* in the NAc core. There was no significant difference in *V*max between adult and adolescent rats (*t*_18_=0.863, *p*=0.399)(Figure 1D). Probability of dopamine release is another dopamine dynamic that can impact stimulated dopamine levels (Ferris et al., 2013). We thus compared paired-pulse ratios, a measure of dopamine release probability (Cragg, 2003), between adult and adolescent rats. However, while interstimulus interval impacted paired-pulse ratios, there was no age-related difference (main effect of interstimulus interval: *F*_1.84, 34.97_=28.64, *p*<0.001)(Figure 1E).

**Figure 1.**
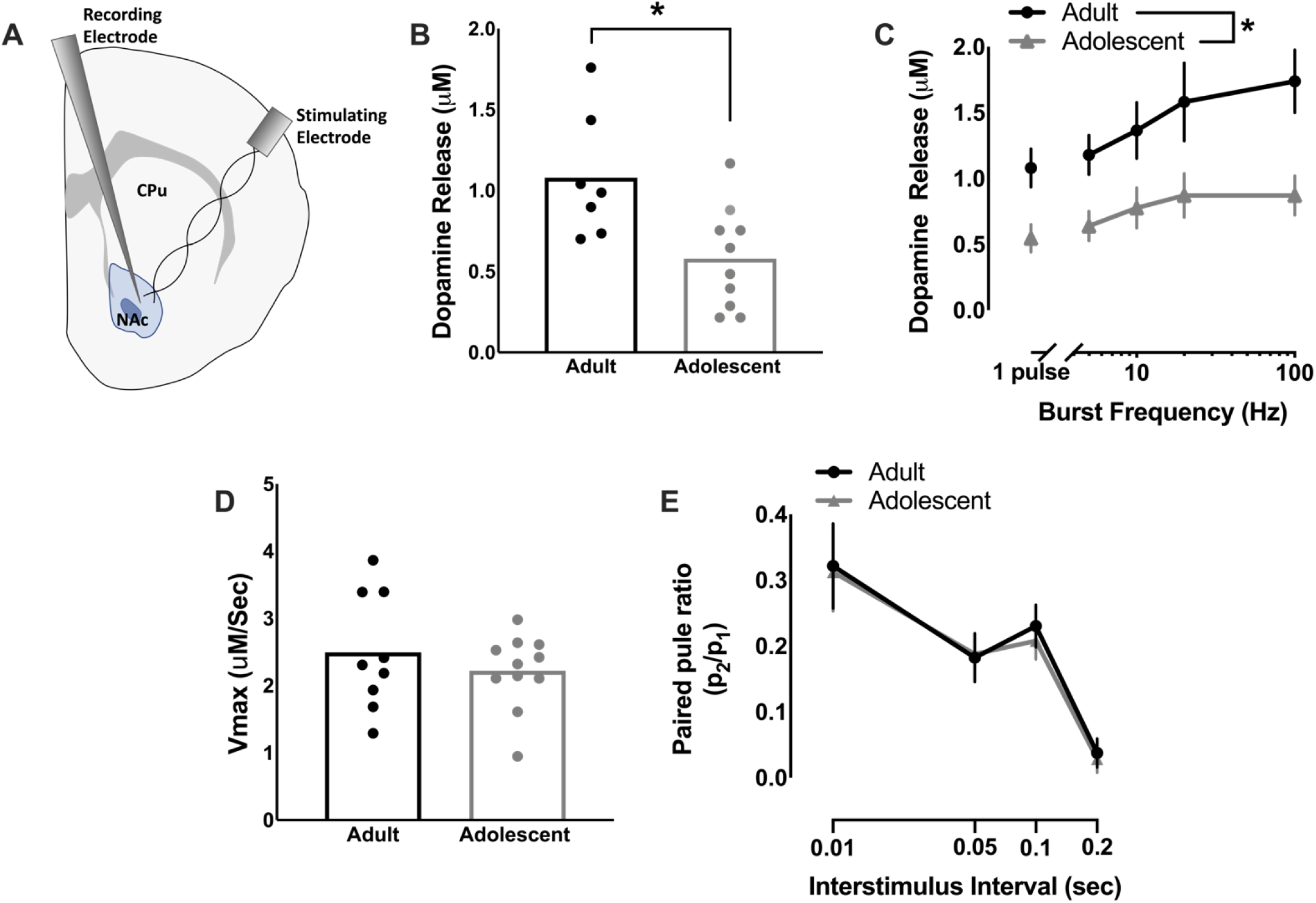
Early adolescent rats have decreased stimulated dopamine release in the nucleus accumbens core. (A) *Ex vivo* fast scan cyclic voltammetry was used to compare dopamine release in the nucleus accumbens core of adult (>P70) and early adolescent (P28-35) male rats. (B) Dopamine release was significantly lower in adolescent than adult rats in the NAc core at single-pulse stimulations and (C) across a range of stimulation parameters that model tonic and phasic firing of dopamine neurons (adults: n=7; adolescents: n=10). (D) Dopamine uptake rates can impact stimulated dopamine release levels (Ferris et al., 2013), so maximal rate of dopamine uptake (*Vmax*) was examined in the NAc core. However, *Vmax* did not differ between adult and adolescent rats (adults: n=9; adolescents: n=11). (E) Differences in probability of dopamine release can also impact stimulated dopamine release (Ferris et al., 2013), so we also used paired-pulse ratios to examine differences in release probability. Paired-pulse ratios did not differ between adult and adolescent rats (adults: n=9; adolescents: n=12). In line graphs, symbols represent means ± SEMs. In bar graphs, bars represent means and symbols represent individual data points. *p < 0.05.

### nAChRs, particularly α6-containing nAChRs, differentially modulate dopamine release in the NAc core of early adolescent rats

Since differences in dopamine dynamics at the dopamine terminal such as uptake or release probability did not appear to mediate the age-related differences in stimulated dopamine release, we then hypothesized that local circuitry modulators of dopamine release may be driving the differences between adult and adolescent rats. Acetylcholine, signaling through multiple sub-types of nAChRs, is an important modulator of dopamine release in the NAc core (Figure 2A). For example, acetylcholine can drive dopamine release independent of action potentials (Threlfell et al., 2012; Cachope et al.,, 2012) and tonically released acetylcholine increases baseline dopamine release (Zhou et al., 2001). Additionally, antagonism or desensitization of nAChRs alters dopamine release in a frequency-dependent manner (Rice and Cragg, 2004). Given the importance of nAChRs as a local mediator of dopamine release, we applied various drugs that selectively targeted each sub-type common in the NAc core (Livingstone and Wonnacott, 2009) and compared changes in dopamine release in adults and adolescents. MLA, a selective α7 nAChR antagonist, impacted dopamine release in a frequency-dependent manner, but not in an age-dependent manner (MLA X frequency interaction: *F*_4, 109.654_=4.347, *p*=0.003)(Figure 2B). In contrast, α-Ctx, a selective antagonist of α6-containing nAChRs, differentially impacted dopamine release in adult and adolescent rats (age X α-Ctx interaction: *F*_1, 161.896_=27.218, *p*<0.001), decreasing dopamine release to tonic-like stimulations in adults (1 pulse: *t*_6_=5.743, *p*=0.0012; 5 pulse 5Hz: *t*_6_=6.917, *p*<0.001; 5 pulse 10Hz: *t*_6_=4.927, *p*=0.0026) and increasing dopamine release to phasic-like stimulation in adolescents (5 pulse 20Hz: *t*_10_=2.335, *p*=0.0417; 5 pulse 100Hz: *t*_10_=3.276, *p*=0.0084)(Figure 2C). When DHβE, a β2-selective antagonist, was applied following α-Ctx, it further decreased dopamine release, but did not impact dopamine release differently by age group (main effect of DHβE: *F*_1, 161.022_=11.787, *p*=0.001)(Figure 2F). Interestingly, when we compared dopamine release and the impact of α-Ctx on dopamine release in mid-adolescent rats (P39-42) and adults (>P70), there was no age-related difference in raw dopamine release (main effects of age and age X frequency interaction: *F*<1)(Figure 3A) or in α-Ctx modulation of dopamine release in the NAc core (main effect of drug: *F*_1, 80.984_=64.613, *p*<0001)(Figure 3B).

**Figure 2.**
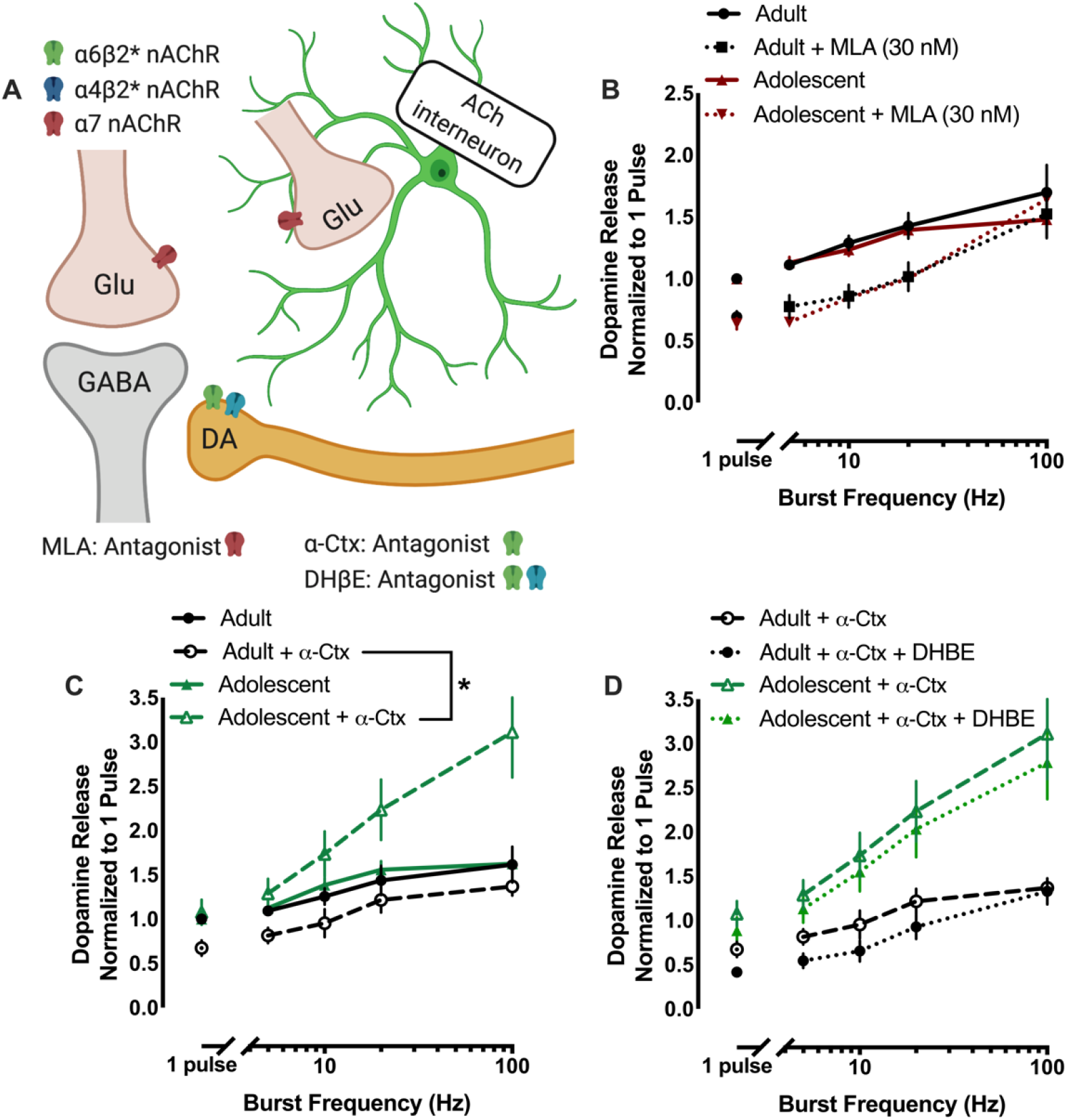
Nicotinic acetylcholine receptors (nAChRs), and in particular α6-containing nAChRs, differentially regulate dopamine release in the nucleus accumbens core of adolescent rats. (A) Acetylcholine, signaling through nAChRs located on dopamine terminals, is an important regulators of local dopamine release (Rice and Cragg, 2004). There are three main types of nAChRs found in the nucleus accumbens core: α7, α6β2-containing, and α4β2-containing. A schematic shows the localization of the sub-types of nAChRs in the nucleus accumbens core and lists the action of drugs bath applied to brain slices to examine nAChR mediation of dopamine release. We used antagonists specific to various sub-types of nAChRs to examine which nAChRs were playing a role in mediating the age-related difference in nAChR modulation of dopamine release. (B) MLA, a selective α7 nAChR antagonist, decreased dopamine release, but did not differentially effect dopamine release in adult and adolescent rats (adults: n=8; adolescents: n=7). (C) In contrast, α-Ctx, an α6-containing nAChR selective antagonist, facilitated dopamine release in adolescents, but decreased dopamine release in adult rats (adults: n=8; adolescents: n=12). (D) Further antagonism of non-α6 β2-containing nAChRs, using DHβE, additionally decreased dopamine release, but not in an age-specific manner (adults: n=8; adolescents: n=12). Symbols represent means ± SEMs. *p < 0.05.

**Figure 3.**
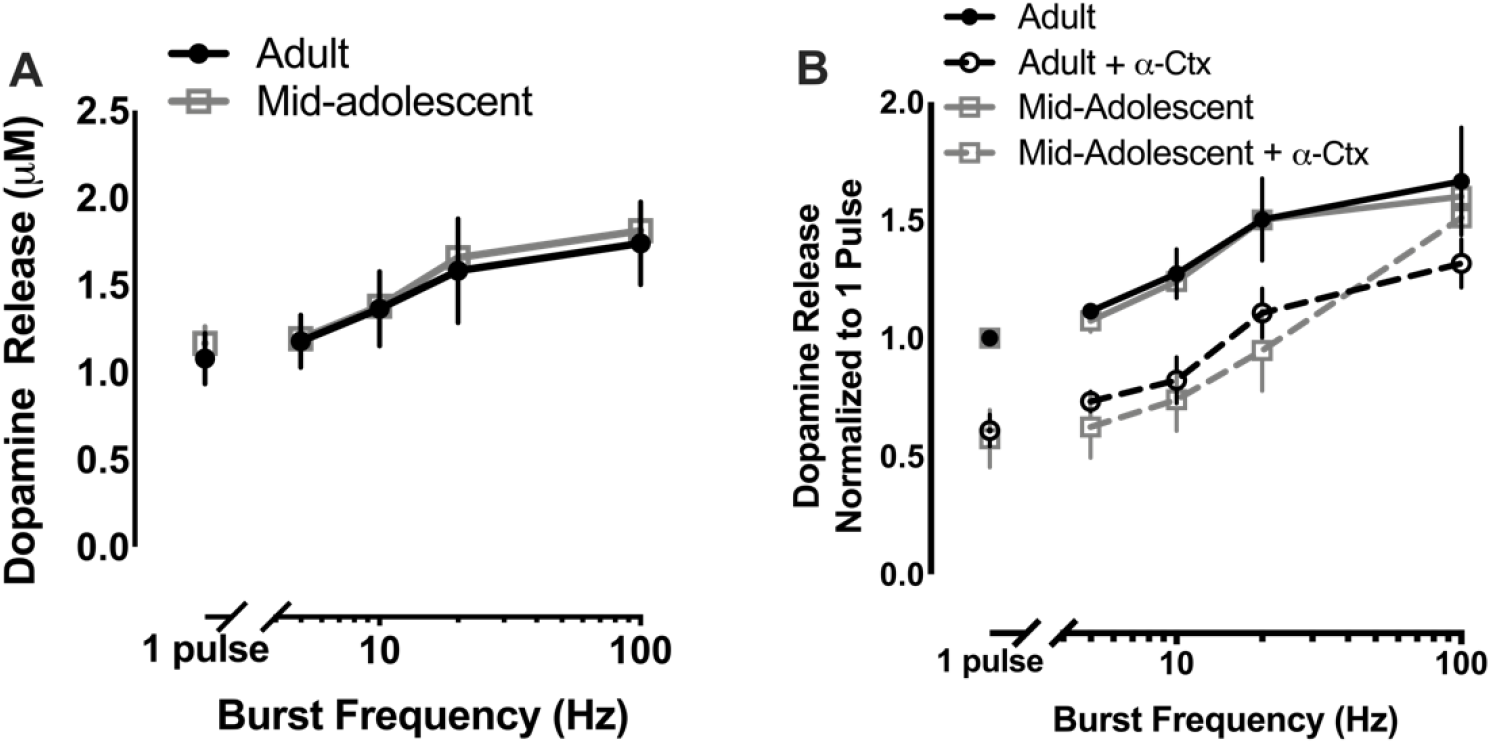
Dopamine release and α6-containing nicotinic acetylcholine receptor modulation of dopamine release do not differ between mid-adolescence and adulthood in rats. (A) Stimulated dopamine release in the nucleus accumbens core did not differ between adult (>P70) and mid-adolescent (P39-42) male rats. (B) α-Ctx, a selective α6-containing nAChR antagonist, decreased dopamine release to a similar degree in adult and mid-adolescent rats (adults: n=7; mid-adolescents: n=4 for A and B). Symbols represent means ± SEMs. *p < 0.05.

### Differences in modulation of dopamine release by α-Ctx in the adolescent NAc core are mediated by GABA

α6-containing nAChRs are positioned on the terminal of dopamine neurons in the NAc core (see Yang et al., 2009). To determine whether α-Ctx is differentially impacting terminal dopamine dynamics in an age-specific manner, we examined *V*max and paired-pulse ratios. α-Ctx did not impact *V*max differently in adult and early adolescent rats (*t*_17_=0.131, *p*=0.898)(Figure 4A). Additionally, α-Ctx did increase paired-pulse ratios with the greatest increase at the smallest interstimulus interval, indicating a decreased likelihood of dopamine release (α-Ctx X insterstimulus interval interaction: *F*_3,119.685_=2.978, *p*=0.034). However, this effect was not different between ages (age X α-Ctx interaction: *F*_1, 120.561_=0.047, *p*=0.829)(Figure 4B).

**Figure 4.**
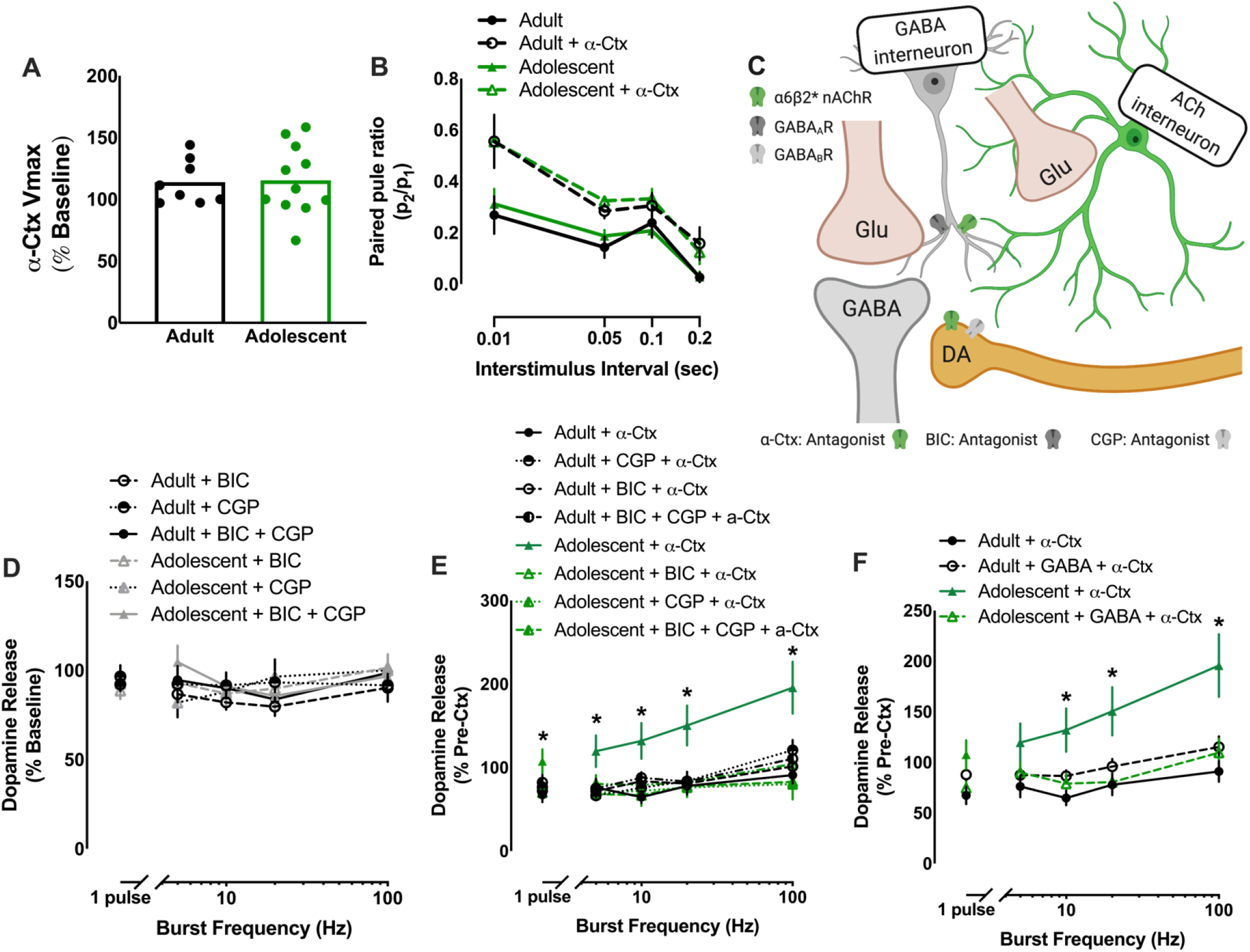
Differential modulation of dopamine release by α6-containing nicotinic acetylcholine receptors in adolescent rats is mediated through GABA receptor signaling. (A) Given the differential effect of antagonizing α6-containing nicotinic acetylcholine receptors (nAChRs) on adolescent and adult dopamine release, we examined whether α-Ctx, a selective α6-containing nAChR antagonist, changed dopamine dynamics in an age-specific manner. Maximal rate of dopamine uptake (*Vmax)* was not differentially altered by α-Ctx in adult and adolescent rats (adults: n=8; adolescents: n=11). (B) Release probability of dopamine, measured by paired-pulse ratio, was decreased by α-Ctx, but not in an age-specific manner (adults: n=8; adolescents: n=12). (C) We next hypothesized that the age-related differences in α6-containing nAChR-modulation of dopamine release may be mediated by α-Ctx causing reduction of GABA signaling in adolescents. GABA receptors regulate dopamine release (Brodnik et al., 2019) and nAChRs are found on GABA interneurons (Tepper et al., 2018). A schematic shows potential localization of nAChRs and GABA receptors in the nucleus accumbens core. (D) Bath application of bicuculline (BIC; a selective GABA_A_ receptor antagonist) and CGP-52432 (CGP; a selective GABA_B_ receptor antagonist) alone or together did not impact dopamine release (adults: n=6-8; adolescents: n=5-7). (E) However, prior application of BIC and/or GCP completely blocks the faciliatory effect of α-Ctx on dopamine release in adolescent rats. Antagonism of GABA receptors did not impact the effect of α-Ctx on dopamine release in adult rats (adults: n=6-8; adolescents: n=5-12). (F) We also hypothesized that application of exogenous GABA would block the faciliatory effects of α-Ctx in adolescents by replacing the endogenous GABA reduced by α-Ctx. As hypothesized, prior application of GABA did not impact the effects of α-Ctx in adults, but blocked the α-Ctx-induced increase in dopamine in adolescents (adults: n=6-8; adolescents: n=6-12). In line graphs, symbols represent means ± SEMs. In bar graphs, bars represent means and symbols represent individual data points. *p < 0.05.

Since antagonism of α6-containing receptors did not impact terminal dopamine dynamics in an age-specific manner, we next hypothesized that age-related differences in the impact of α-Ctx may be driven through another local circuitry modulator of dopamine release. Bath application of α-Ctx increases dopamine release in adolescents, indicating that blocking α6 nAChRs leads to disinhibition of dopamine release in adolescents, but not adults. GABA has been shown to inhibit terminal dopamine release (Pitman et al., 2014; Melchior et al., 2015; Brodnik et al., 2019; Lopes et al., 2019) indicating that α-Ctx could be reducing GABA tone in adolescents. In addition, some striatal GABA interneurons express nAChRs (see Tepper et al., 2018)(Figure 4C). To determine whether the differential modulation of dopamine release by α-Ctx in early adolescent and adult rats is mediated through a multi-synaptic mechanism mediated by GABA, we antagonized GABA_A_ and GABA_B_ receptors, alone or together, prior to the bath application of α-Ctx and then compared the effects of α-Ctx in the presence or absence of GABA receptor antagonism. If the age-related effects of α-Ctx are being mediated through GABA receptors, then antagonism of GABA receptors prior to α-Ctx application should eliminate differences in dopamine release modulation between adult and adolescent rats. Application of GABA_A_ and/or GABA_B_ receptor antagonists did not impact dopamine release and did not modulate dopamine release differently in adult and adolescent rats (main effect of age: *F*_1, 26.121_=0.001, *p*=0.981)(Figure 4D). Importantly, however, application of GABA_A_ or GABA_B_ antagonists, alone or together, prior to the application of α-Ctx completely blocked the differential effect of antagonizing α6-containing nAChRs on dopamine release in adult and adolescent rats (age X frequency X drug interaction: *F*_12, 193.305_=3.988, *p*<0.001)(Figure 4E). To further test the hypothesis that GABA is mediating the differential effects of α-Ctx in adolescents and adults, we bath applied exogenous GABA prior to the application of α-Ctx. If α-Ctx disinhibits dopamine release in adolescents by reducing endogenous release of GABA, then increasing exogenous GABA should by pass the GABA inhibition and prevent the age-related effects of α-Ctx in adolescents. Supporting this, we found that GABA blocks the α-Ctx-induced increase in dopamine release in adolescents, causing adolescents to respond to α-Ctx similarly to adult rats (age X frequency X drug interaction: *F*_4, 110.026_=3.189, *p*=0.016)(Figure 4F).

## Discussion

In the present report, we utilized *ex vivo* FSCV to compare stimulated dopamine release and its regulation in the NAc core of adult and adolescent male rats. Replicating our previous findings (Pitts et al., 2020), we see decreased stimulated dopamine release in the NAc core of male early adolescent rats. We also found this decrease in stimulated dopamine release is not driven by differences in release probability or transporter function (i.e. homoosynaptic mechanisms). Instead, we found that there is multisynaptic regulation of dopamine release by GABA and α6-containing nAChRs that is unique to early adolescent rats. The differences in baseline dopamine release and α6-containing nAChRs modulation of dopamine release found in early adolescents is absent in slightly older mid-adolescent animals (P39-42), who instead have dopamine dynamics that match those of adult rats.

Acetylcholine and nAChRs are important regulators of terminal dopamine release in the striatum (see Cachope and Cheer, 2014; Collins and Saunders, 2020). For example, activation of CINs can cause dopamine release independent of action potentials (Cachope et al., 2012), while pauses in CINs during reward presentations increases dopamine release to burst firing of VTA neurons (Cragg, 2006). In concurrence with this, antagonism of nAChRs has been shown to regulate dopamine release in a frequency-dependent manner, decreasing dopamine release to tonic-like stimulations, while not impacting or facilitating dopamine release to phasic-like stimulations (Rice and Cragg, 2004; Cragg, 2006). We found differences between adult and adolescent rats in modulation of NAc core dopamine release by α6-containing nAChRs, but not by α7 or non-α6 β2* nAChRs. Additionally, the application of α-Ctx (a specific α6-containing nAChRs antagonist) had a completely different pattern of effect on dopamine release in adults than early adolescent rats: decreasing dopamine release to tonic-like firing in adults, while increasing dopamine release to stimulation frequencies modeling burst firing in adolescents. A previous study from our lab found a similar pattern in how α6-containing nAChRs modulated dopamine release in a different model of populations vulnerable to SUDs (the high and low responder model of sensation seeking) (Siciliano et al., 2017). This is an intriguing finding and may indicate that α6-containing nAChR regulation of dopamine is a biomarker for vulnerability to SUD, although additional experiments and studies across more models of vulnerability are necessary to strengthen this conclusion.

α6-containing nAChRs are highly expressed on presynaptic terminals in the mesocorticolimbic dopamine system (see Yang et al., 2009). Antagonism of α6-containing nAChRs causes frequency-dependent regulation of dopamine release, which is hypothesized to occur by altering Ca^2+^ entry and decreasing probability of neurotransmitter release (Soliakov and Wonnacott, 1996; Siciliano et al., 2017). As shown previously (Exley et al., 2008), antagonism of α6-containing nAChRs decreased probability of dopamine release. However, we showed that α-Ctx did not change release probability in an age-dependent manner. Therefore, α6-mediated changes in release probability of dopamine release cannot underlie the effects of α-Ctx in both adolescent and adult rats. Instead, we found that antagonizing GABA_A_ and/or GABA_B_ receptors prior to administration of α-Ctx blocked α-Ctx-induced facilitation of phasic-like dopamine release in adolescents, but did not alter the effects of α-Ctx in adults. This suggests that blockade of α6-containing nAChRs changes homosynaptic regulation of terminal dopamine release in adults, whereas a unique multisynaptic mechanism of α6-containing nAChRs reducing GABA modulation of dopamine offsets the direct terminal effects of α-Ctx and drive increased dopamine release in adolescents. This is further supported by our finding that exogenous replacement of GABA causes α-Ctx to impact dopamine release the same in adults and adolescents, likely through direct effects at the dopamine terminal. Although prior research has found no difference in the density of α6-containing nAChRs in the NAc of early adolescents and adults (Doura et al., 2008), differences in localization or functionality of either α6-containing nAChRs or GABA receptors may underlie adolescent differences that create this novel multisynaptic mechanism of dopamine regulation in adolescents.

nAChRs are hypothesized to be expressed on the pre-synaptic terminals of GABA interneurons in the striatum (see Tepper et al., 2018) and functional evidence indicates GABA_A_ (and possibly GABA_B_) receptors are located on dopamine terminals and can impact dopamine release (Pitman et al., 2014; Melchior et al., 2015; Brodnik et al., 2019; Lopes et al., 2019). Antagonizing GABA_A_ or GABA_B_ receptors, alone or together, did not impact dopamine release. This is consistent with prior literature showing GABA antagonism has no impact on dopamine release (Pitman et al., 2014; Brodnik et al., 2019). Interestingly, one report found that antagonizing GABA_B_ receptors increased dopamine release when β2-containing nAChRs were antagonized (Lopes et al., 2019), indicating that GABA and nAChRs have a potentially complex and reciprocal regulation of dopamine release.

It was unexpected that antagonizing either GABA_A_ or GABA_B_ receptors both blocked the effects α-Ctx in adolescents and that blocking both receptors did not further change the effects of α-Ctx. However, a previous study found that GABA_A_ receptor regulation of dopamine release in the NAc core functions through GABA_B_ receptors (Brodnik et al., 2019). It is possible that a similar mechanism of action is mediating the effects seen here, although further studies are needed to understand the microcircuitry mediating the effect of α6-containing nAChR-regulation of dopamine release in adolescent rats.

In summary, stimulated dopamine release is decreased in the NAc core of early adolescent male rats. This effect is potentially driven, in part, by multisynaptic regulation of dopamine release in adolescent rats, through GABA and a6-containing nAChRs, that is not present in adult rats. The dopamine system undergoes reorganization during adolescence (see Wahlstrom et al., 2010a,b; Padmanabhan and Luna, 2014) and these changes are implicated in behavioral changes and increased vulnerability to the development of psychiatric disorders (Spear, 2000, 2013; Nelson et al., 2005; Wahlstrom et al., 2010a,b). These changes also increase vulnerability to the development of long-lasting psychiatric disorders, particularly following perturbations that may disrupt brain development occurring during adolescence. For example, preclinical studies find that stress (such as social isolation) or drug use in adolescence can alter decision making, impair cognition, and increase drug use in adulthood, even when stress or drug use are constrained to adolescence (Butler et al., 2016; Spear, 2016; DePoy et al., 2017; Hinton et al., 2019). This study gives us novel insight into age-related differences in dopamine release and its regulation. Understanding how these systems develop in healthy individuals is imperative, so we can understand how perturbations may disrupt development, leading to disease states.

## Acknowledgments

Special thanks to Miss Lacey Sexton for her contributions. This work was supported by the National Institutes of Health grants R00 DA031791 (MJF), P50 DA006634 (MJF), P50 AA026117 (MJF), K12 GM102773 (EGP), F32 AA028162 (EGP), and the Peter McManus Charitable Trust.

## Author Contributions

Conceptualization, E.G.P and M.J.F.; Formal analysis, E.G.P.; Investigation, E.G.P.; Resources, M.J.F.; Writing ‐ Original Draft, E.G.P and T.A.S.; Writing ‐ Review & Editing, E.G.P., T.A.S., and M.J.F.; Funding Acquisition, E.G.P and M.J.F.

## STAR Methods

### Animals

Early adolescent (P28-35), mid-adolescent (P38-42) and adult (P70-90) male Sprague-Dawley rats (Envigo, Huntingdon, UK) were maintained on a 12:12 h reverse light/dark cycle (4:00 a.m. lights off; 4:00 p.m. lights on) with food and water available *ad libitum*. All animals were maintained according to the National Institutes of Health guidelines in Association for Assessment and Accreditation of Laboratory Animal Care accredited facilities. All experimental protocols were approved by the Institutional Animal Care and Use Committee at Wake Forest School of Medicine.

### Slice preparation

Rats were anesthetized with isoflurane and then euthanized by rapid decapitation in a ventilated area free of blood or tissue from previous animals. Brains were rapidly removed and transferred into pre-oxygenated (95% O_2_ / 5% CO_2_) artificial cerebral spinal fluid (aCSF) containing (in mM): NaCl (126), KCl (2.5), monobasic NaH_2_PO_4_ (1.2), CaCl_2_ (2.4), MgCl_2_ (1.2), NaHCO_3_ (25), dextrose (D-glucose) (11), and L-ascorbic acid (0.4). Tissue was sectioned into 400 mm-thick coronal slices on a compresstome^®^ VF-300 vibrating microtome (Precisionary Instruments Inc., San Jose, CA). Brain slices were transferred to testing chambers containing oxygenated aCSF (32 °C) flowing at 1 mL/min.

### Ex vivo fast scan cyclic voltammetry

*Ex vivo* FSCV provides unique advantages for studying local circuit modulation of dopamine release. For example, FSCV has high spatial and temporal resolution and allows the examination of uptake kinetics and dopamine release while modeling of a wide range of neuronal firing patterns (see Ferris et al., 2013). Here, *ex vivo* FSCV was used to characterize dopamine release in the NAc core (Fennell et al., 2020). Briefly, a <u>carbon-fiber</u>recording microelectrode was placed 100-150 mM from a bipolar stimulating electrode. Dopamine release was initially evoked by a single electrical pulse (750 mA, 2 msec, monophasic) applied to the tissue every 3 minutes. After the extracellular dopamine response was stable (3 collections within <10% variability), five-pulse stimulations were applied at varying frequencies (5, 10, 20, or 100 Hz) to model the physiological range of dopamine neuron firing. Additionally, paired-pulse stimulations were used to determine baseline probability of dopamine release. Paired-pulse ratios can be used to determine the release probability of dopamine (Condon et a., 2019; Cragg et al., 2003). We define paired-pulse ratio as the ratio P2/P1, where P1 is peak DA release detected following 1p stimulus and P2 is the peak DA release attributable to the second stimulation only. P2 was determined by subtracting the DA release, including decay phase, after a single pulse from the summed paired-pulse response.

After assessing the dopamine response to varying stimulation parameters, various compounds targeting nAChRs (Methyllycaconitine [MLA; a selective α7 nAChR antagonist], 30 nM; α-conotoxin PIA [α-Ctx; a selective α6-containing nAChR antagonist], 100 nM; dihydro-beta-erythroidine [DhβE; a selective β2-containing nAChR antagonist], 500 nM) or GABARs (Bicuculline [BIC; a selective GABA_A_R antagonist], 10 μM; CGP-52432 [CGP; A selective GABA_B_R antagonist], 5 μM; γ-Aminobutyric acid [GABA], 10 μM) were bath applied and dopamine response equilibrated to single pulse stimulation. Separate slices from the same animal were used to test each drug independently, and the same frequency-response curves assessed under drug-free conditions were reassessed following drug application in each slice. In order to test the distinct contributions of α6* and non-α6* nAChRs, we added α-Ctx and DhβE in a cumulative fashion, equilibrating and testing single and multi-pulse frequencies (described above) following α-Ctx and then DhβE. Changes in dopamine signaling between α-Ctx alone and in combination with DHβE differentiated the contribution of α6* and non-α6* β2-containing nAChRs. Similarly, to examine the role of GABA or GABARs in the effects of α-Ctx, we bath applied GABA or BIC and CGP alone or together. Then, after equilibration and frequency-response curves, we applied α-Ctx and re-tested single- and multi-pulse stimulations.

### Drugs

Methyllycaconitine citrate (20-ethyl-1α,6β,14α,16β-tetramethoxy-4-[[[2-[(3S)-3-methyl-2,5-dioxo-1-pyrrolidinyl]benzoyl]oxy]methyl]-aconitane-7,8-diol, 2-hydroxy-1,2,3-propanetricarboxylate; Cayman Chemical Company, Ann Arbor, MI), α-conotoxin PIA (Research and Diagnostic Systems, Inc., Minneapolis, MN), Dihydro-β-erythroidine hydrobromide ((2*S*,13b*S*)-2-Methoxy-2,3,5,6,8,9,10,13-octahydro-1*H*,12*H*-benzo[*i*]pyrano[3,4-*g*]indolizin-12-one hydrobromide; Tocris Bioscience, Bristol, UK), (-)-Bicuculline methochloride ([*R*-(*R*\*,*S*\*)]-5-(6,8-Dihydro-8-oxofuro[3,4-*e*]-1,3-benzodioxol-6-yl)-5,6,7,8-tetrahydro-6,6-dimethyl-1,3-dioxolo[4,5-*g*]isoquinolinium chloride; Tocris Bioscience, Bristol, UK), CGP-52432 (3-[[(3,4-Dichlorophenyl)methyl]amino]propyl] diethoxymethyl)phosphinic acid; Tocris Bioscience, Bristol, UK), and γ-Aminobutyric acid (Tocris Bioscience, Bristol, UK) were dissolved in distilled water. 1 mM concentration aliquots were stored at -20°C or 4°C and diluted with oxygenated aCSF to final concentration before bath application on slices.

### Data Analysis

Demon Voltammetry and Analysis software was used to acquire and model FSCV data (Yorganson et al., 2011). Recording electrodes were calibrated by recording electrical current responses (in nA) to a known concentration of dopamine (3 mM) using a flow-injection system. This was used to convert electrical current to dopamine concentration. Michaelis-Menten kinetics were used to determine maximal rate of dopamine uptake (*Vmax*) (Ferris et al., 2013).

### Statistical Analysis

Sample size estimation for all experiments was based on effect size and standard deviations from prior published research using voltammetry in order to detect small-to medium-sized effects. Baseline dopamine release to single-pulse stimulations and *Vmax* (raw numbers and percent change) were compared by students t-test. Baseline dopamine release to multi-pulse stimulations and paired-pulse ratio were compared by two-way mixed-factor ANOVA. Dopamine release following drug application (including percent change, normalized to 1 pulse pre-drug baseline, and paired-pulse ratio) were compared by three-factor generalized linear mixed model analysis. In the case of significant interactions, Bonferroni or Tukey post-hoc comparisons were used. Graph Pad Prism (version 8, La Jolla, CA) or SPSS (version 24, International Business Machine Corporation, Armonk, NY) were used to statistically analyze data sets (with α δ 0.05) and compose graphs. Values >2 standard deviations above or below the mean were considered outliers and excluded. Data are presented as mean ± SEM for group data across multiple variables or mean with individual data points for bar graphs.

